## supplementary for "Neuronal integration in the adult olfactory bulb is a non-selective addition process"

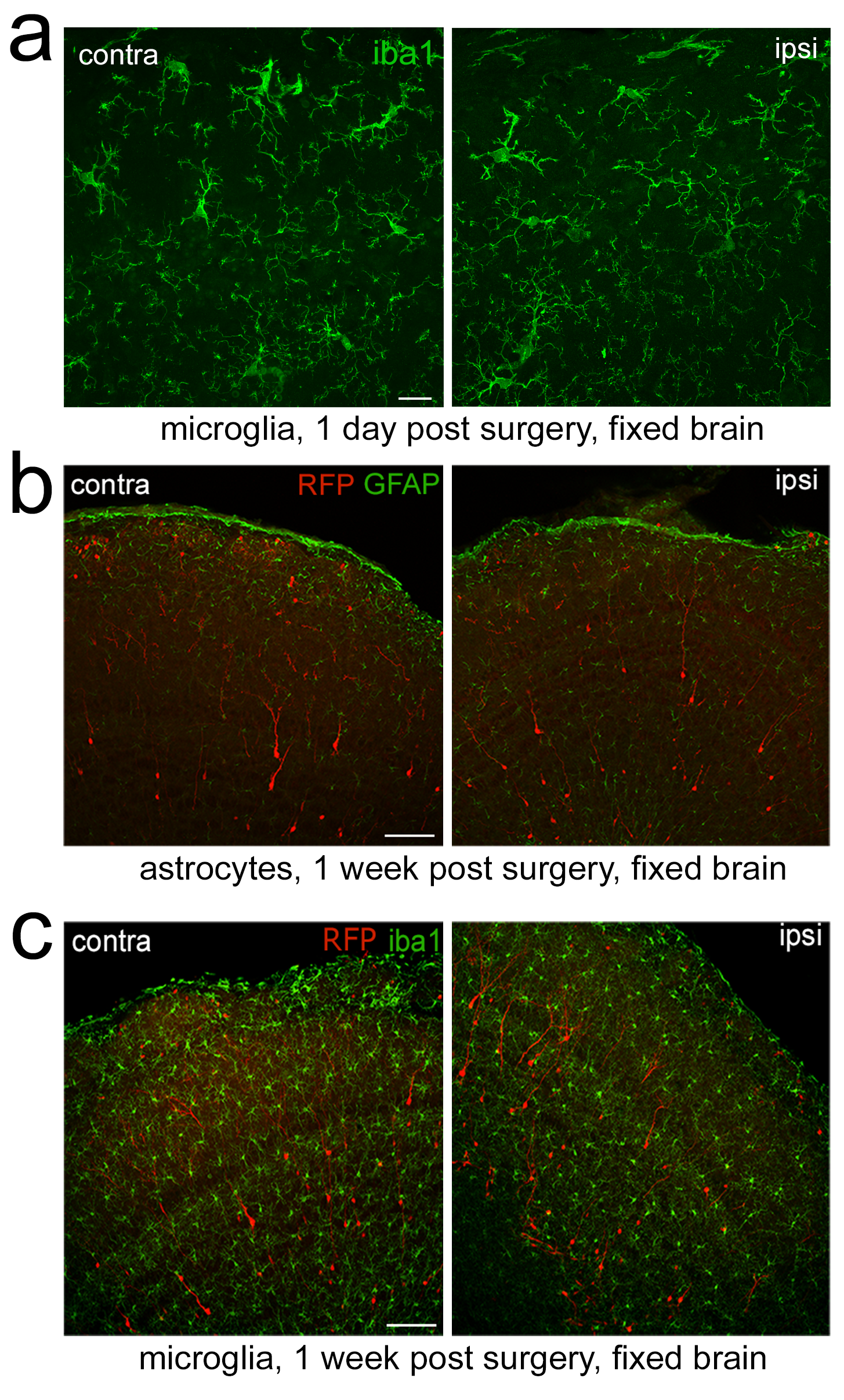

Supplementary Fig. 1

a. Immunostaining for the microglial marker iba1 demonstrates the absence of a detectable microglia reaction at 1-day post surgery.

b. Comparison of GFAP staining in the OB. Control and window carrying OB are indistinguishable at 1-week post surgery.

c. Immunostaining for the microglial marker iba1 demonstrates the absence of a detectable microglia reaction at 1-week post surgery. Scale bar: 10 μm in a, 50 μm in b,c.

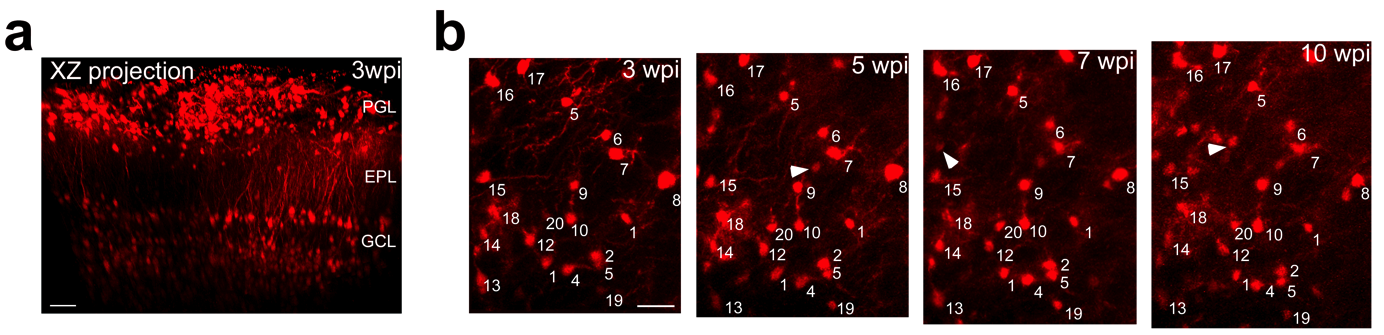

Supplementary Fig. 2

A. XZ projection of the OB showing that the PGL, EPL and the superficial GCL can be reached through a thin skull preparation. b. Example of Z-projection identifying individual granule neurons between 3 and 10 weeks post induction in perinatal mice. The first cohort is numbered. Additional neurons of later cohorts appeared in the observation window over successive imaging sessions (arrowheads). Scale bar: 50 μm in a, 30 μm b.

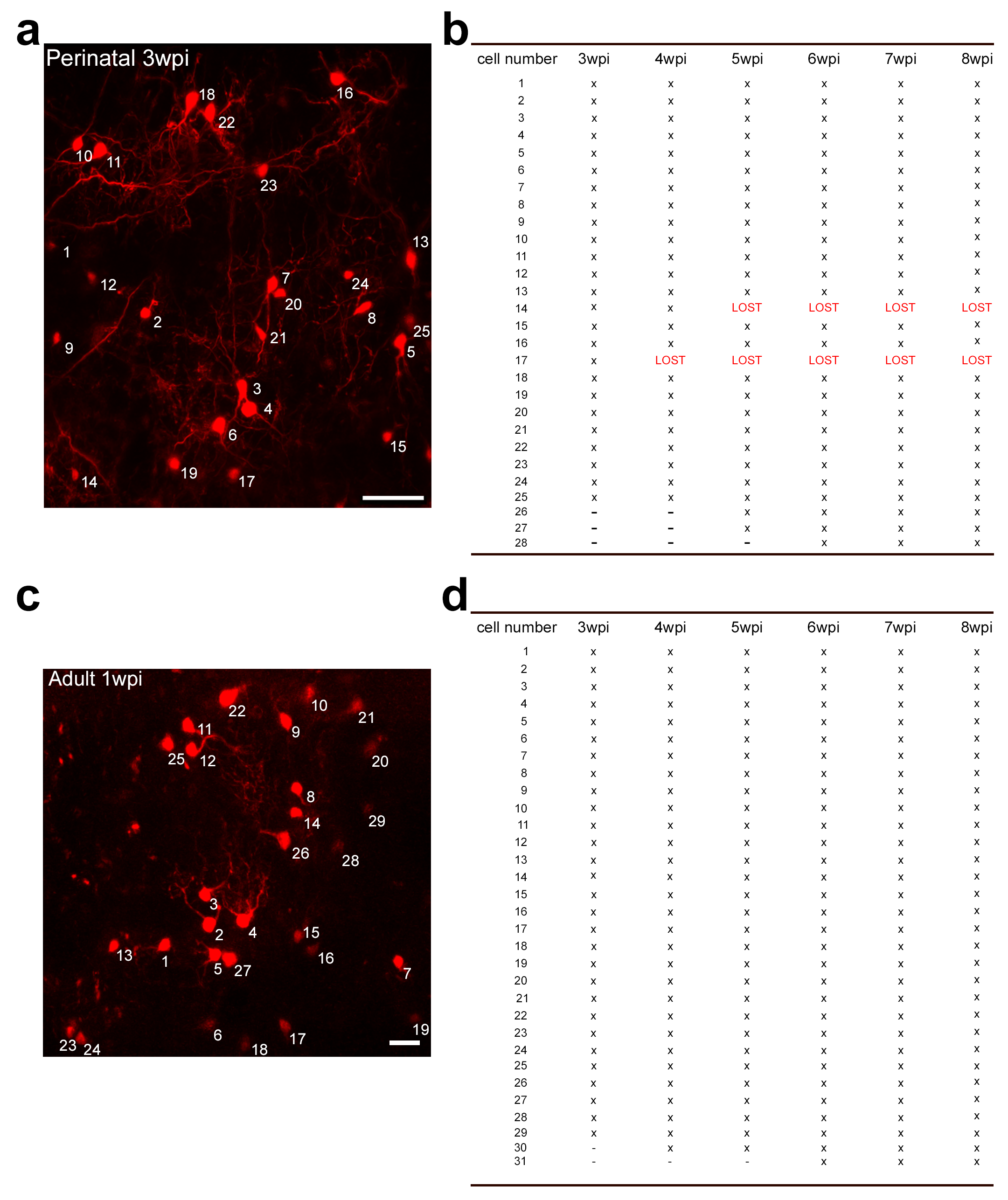

Supplementary Fig. 3

Image and table exemplifying how perinatally (a,b) and adult (c,d) generated neurons were scored over the critical period (for a,b compare also Fig. 1d). Scale bar: 50 μm in a, 20 μm b.

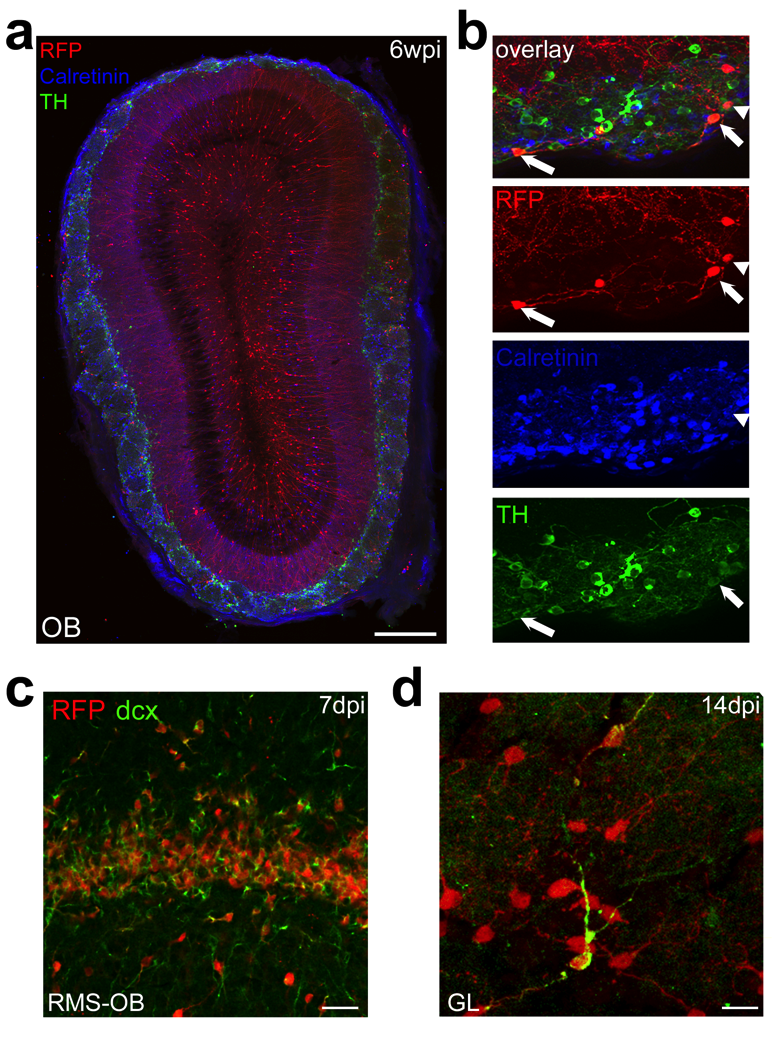

Supplementary Fig. 4

a, b. Coronal section through the OB of an adult Nestin-Cre-ERT2/Rosa-RFP mouse 6 weeks after tamoxifen injection at 2 months of age stained for the OB neuron subtype markers TH and Calretinin.

c, d. Seven days post induction (dpi) large amounts of OB neurons in the OB express the immature neuronal marker doublecortin (dcx). At 14 dpi individual cells in the GCL still express dcx. Scale bar: 300 μm in a, 30 μm in c, 10 μm in d.

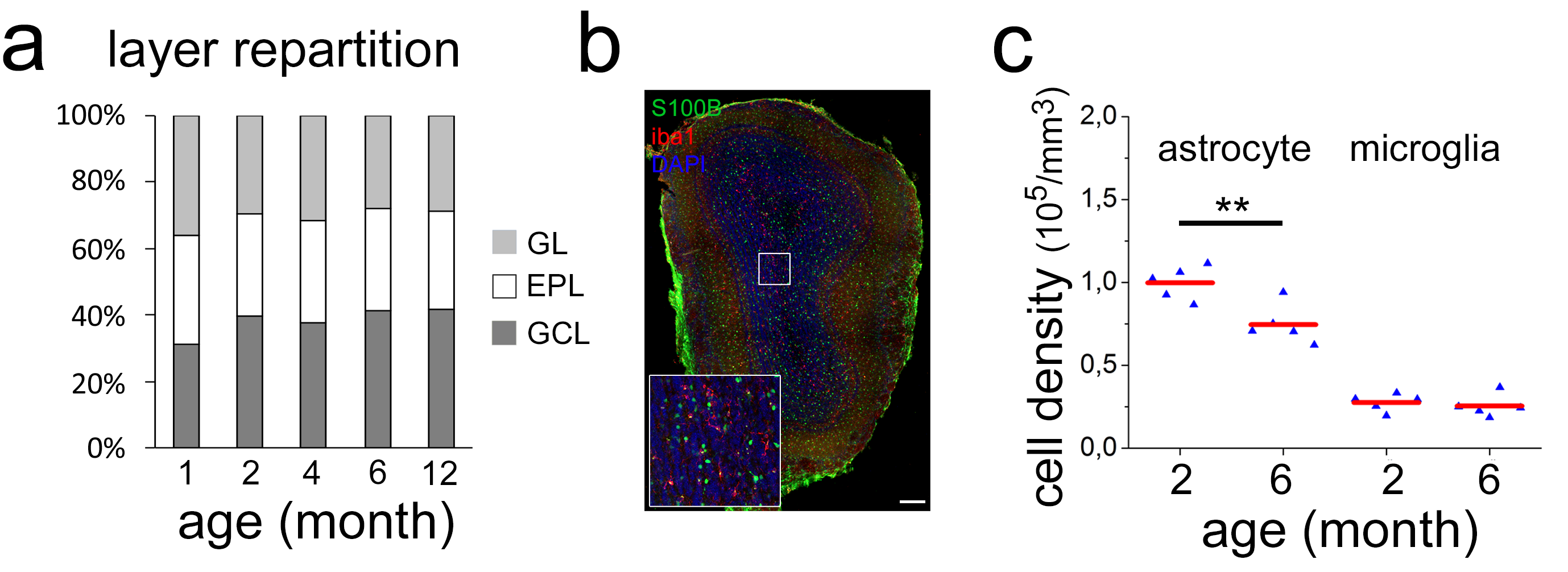

Supplementary Fig. 5

**a.** Repartition of OB layers (GL, EPL, GCL) with increasing age. Although the volume of all layers increases with time their ratio remains constant.

**b.** Coronal section through the OB of a 6 month old C57Bl6 stained for the astrocytic marker S100b, the microglial marker iba1 and the nucleus with DAPI.

**c.** Quantification of astrocytic and microglial density between 2 and 6 months. Astrocyte density is significantly decreased (p=0,006) while microglia density is constant. Scale bar: 200 μm in b

Supplementary Video 1

Example of a Z-stack showing perinatally born neurons in the GL. This stack was the basis for the projection presented in Fig. 1d.

Supplementary Video 2

Example of a Z-stack showning adult born neurons in the GL. This stack was the basis for the projection presented in Extended Data Fig. 3d.

Supplementary Video 3

3D representation of an adult OB based on light sheet microscopy.
